# Invasion and spread of the neotropical leafhopper *Curtara insularis* (Hemiptera: Cicadellidae) in Africa and North America and the role of high-altitude windborne migration in invasive insects

**DOI:** 10.1101/2024.05.24.595796

**Authors:** Rita Nartey, Lourdes Chamorro, Matt Buffington, Yaw A. Afrane, Abdul R. Mohammed, Christopher M. Owusu-Asenso, Gabriel Akosah-Brempong, Cosmos Manwovor-Anbon Pambit Zong, Solomon V. Hendrix, Adama Dao, Alpha S. Yaro, Moussa Diallo, Zana L. Sanogo, Samake Djibril, Susan E. Halbert, Roland Bamou, Catherine E. Nance, Charles R. Bartlett, Don R. Reynolds, Jason W. Chapman, Kwasi Obiri-Danso, Tovi Lehmann

## Abstract

Invasive insects threaten ecosystem stability, public health, and food security. Documenting newly invasive species and understanding how they reach into new territories, establish populations, and interact with other species remain vitally important. Here, we report on the invasion of the South American leafhopper, *Curtara insularis* into Africa, where it has established populations in Ghana, encroaching inland at least 350 km off the coast. Importantly, 80% of the specimens collected were intercepted between 160 and 190 m above ground. Further, the fraction of this species among all insects collected was also higher at altitude, demonstrating its propensity to engage in high-altitude windborne dispersal. Its aerial densities at altitude translate into millions of migrants/km over a year, representing massive propagule pressure. Given the predominant south-westerly winds, these sightings suggest an introduction of *C. insularis* into at least one of the Gulf of Guinea ports. To assess the contribution of windborne dispersal to its spread in a new territory, we examine records of *C. insularis* range-expansion in the USA. Reported first in 2004 from central Florida, it reached north Florida (Panhandle) by 2008-2011 and subsequently spread across the southeastern and south-central US. Its expansion fits a “diffusion-like” process with 200—300 km long “annual displacement steps”—a pattern consistent with autonomous dispersal rather than vehicular transport. Most “steps” are consistent with common wind trajectories from the nearest documented population, assuming 2—8 hours of wind-assisted flight at altitude. *Curtara insularis* has been intercepted at US ports and on trucks. Thus, it uses multiple dispersal modalities, yet its rapid overland spread is better explained by its massive propagule pressure linked with its high-altitude windborne dispersal. We propose that high-altitude windborne dispersal is common yet under-appreciated in invasive insect species.

## Introduction

Invasive insect species pose extreme threat to biodiversity, ecosystem stability, and human welfare, as many invasives impact public health (e.g., *Aedes aegypti, Yellow Fever virus, Anopheles stephensi*), and food security (e.g., *Ceratitis capitata, Helicoverpa armigera*; (Elton 1958; Hulme et al. 2008; Jones et al. 2019; Li et al. 2020; Lounibos 2002; Mack et al. 2000; Pysek et al. 2020; Sinka et al. 2020; Takken, Lindsay 2019; Turner et al. 2021; WHO 2023). Documenting invasive species and understanding how they reach new territories, establish populations, and interact with other species remain vitally important despite limited success in reversing invasions after establishment of populations in new regions (Anderson et al. 2010; Elton 1958; Hulme et al. 2008; Lehmann et al. 2023; Lounibos 2002; Mack et al. 2000). Typically, dispersal of invasive species is divided into (i) long-range movements, e.g., between continents (ii) long-range spread post-arrival in a new territory, and iii) local short-range spread within and between adjacent habitats. Most attention is focused on cross-continent movements (i) because prevention at this stage would be most effective, yet the ‘secondary spread’ (ii) determines whether the species remains stable near its introduction site (“alien” or “naturalized”) vs. “invasive.” High dispersal capacity is a key trait for invasive species (Hulme et al. 2008; Jones et al. 2019; Mack, Occhipinti 1999; Tsoar et al. 2011). In most insects, these dispersal modalities are believed to be mediated by vehicular transport, especially via the maritime trade (Elton 1958; Hulme et al. 2008; Lounibos 2002; Renault et al. 2018; Turner et al. 2021). However, apart from a few major pests, the dearth of information concerning the majority of introduced insects and inherent bias in our methodologies and expectations may limit understanding of dispersal modalities in invasive species.

Sampling of insects at altitude has often been focused on particular pests such as the desert locusts, the brown planthopper, armyworm moths, and malaria mosquitoes, yet these studies revealed surprising diversity and abundance of insects (Anderson et al. 2010; Chapman et al. 2004; Drake, Reynolds 2012; Florio et al. 2020; Glick 1939; Hu et al. 2016; Huang et al. 2024; Reynolds et al. 1996; Riley, Reynolds 1996; Riley et al. 1995; Wu et al. 2021; Yaro et al. 2022). Many of these windborne-migrant insects have been implicated to cover tens, hundreds and thousands of kilometers in their journey (Chapman et al. 2015; Drake, Reynolds 2012; Fu et al. 2014; Ghauri 1983; Hu et al. 2016; Wu et al. 2021).

The interception of the leafhopper *Curtara insularis* (Caldwell, 1952; Fig. S1) flying at altitude, representing a new continental record for Africa, suggests that high-altitude windborne dispersal plays a key role in the species’ rapid spread post-arrival into the continent. A member of a genus that is endemic to the Western Hemisphere, *C. insularis* was originally known from Argentina, Paraguay, and Brazil (Zahniser, Nartey 2024). It has expanded across the Americas since the early 2000s (Kittelberger et al. 2021; Zahniser, Nartey 2024). Unlike certain members of its family, *C. insularis* is not known to vector plant pathogens or impact any crop. Nonetheless, as it feeds on new host plants in its new range, it might play a new role as a vector of local pathogens. An example of such a case is the glassy-winged sharpshooter (*Homalodisca vitripennis* (Germar)) that has changed transmission patterns of local strains of the plant-pathogenic bacterium, *Xylella fastidiosa*, in its new range, i.e., the Western USA, resulting in epidemics of oleander leaf scorch and Pierce’s disease in southern California (Almeida, Nunney 2015; Pysek et al. 2020).

Typically, the rapid spread of invasive insects is attributed to vehicular transport, which is often involved to some or great extent (below). However, here, we assess the role of high-altitude windborne dispersal in the spread of an invasive insect over a new territory. We present results from our aerial and ground-level surveillance in Africa as well as an analysis of a new compilation of distributional records of *C. insularis* in the US based on multiple data sources including citizen-science databases. We offer a descriptive, semiquantitative framework to ascertain the relative contribution of windborne spread versus vehicular spread using spatio-temporal records and data on wind patterns. Based on our results, we propose that high-altitude windborne dispersal is especially common in many invasive insect species, in the hope it would be subject to a rigorous test in the near future.

## Methods

### Sampling Sites

Aerial collection stations were established in rural open areas in Ghana and Mali (Fig. 1A). In southern Ghana two stations were set up in the moist-semi-deciduous forest near the town of Agogo (N:6.961, W: −0.960), and in the Guinea woodland ecozone near the town of Wenchi (N:7.781, W:-2.162). In southern Mali one station was set in the Sudano-Guinean ecozone near the village Kenieroba (N:12.112, W:-8.332). Agogo, Wenchi, and Kenieroba receive 1200-1400, 1000-1200, and 900-1000 mm of rain annually, respectively.

**Fig. 1.**
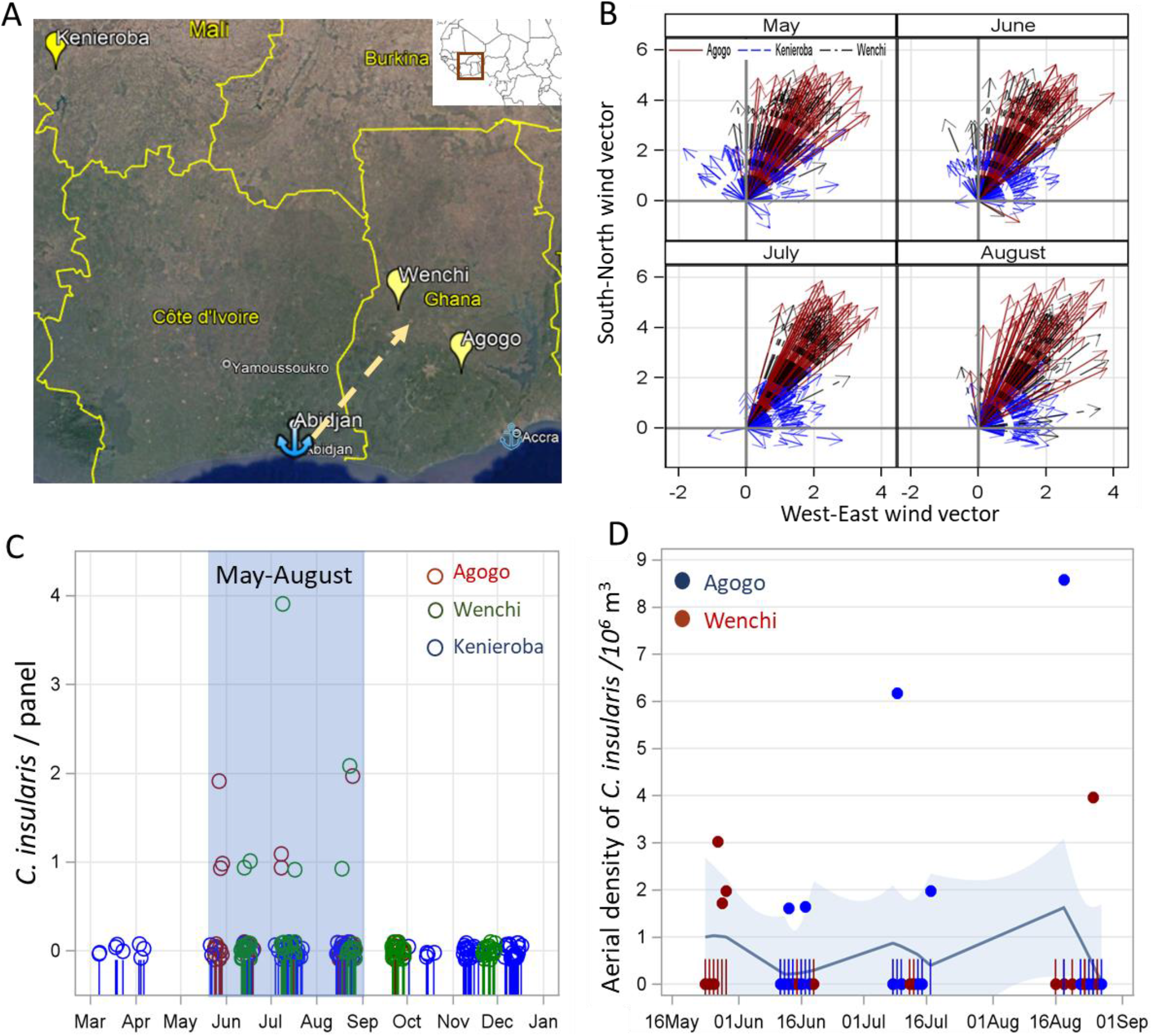
Study area showing collection sites with wind patterns (during inferred migration season) and panel and aerial densities of *C. insularis*. A) Map showing sampling sites in Ghana and Mali (balloon symbols) with the port of Abidjan, Ivory Coast (blue anchor). The arrow shows the predominant wind direction. B) Nightly wind at altitude (300 m agl) by month during 2020-2021 at each sampling station (color) showing the direction (arrow) the insects will be carried towards, from the aerial sampling site (origin). The wind speed is indicated by the vector length (source: ERA5). C) Panel density of *C. insularis* at the different sampling stations (regardless of altitude) with corresponding color fringe (bottom) indicates sampling night used in this study. Blue shade indicates period of interception of *C. insularis*. D) Aerial densities of *C. insularis* at Agogo and Wenchi at altitude (based on panels at 160-190 m agl). Fringe (bottom) indicates sampling dates at altitude in Ghana.

However, in Kenieroba, the rainy season is confined to May-November whereas the Ghanian sites receive rains most months of the year (Siebert 2014). These areas are interspersed with farmland along rivers and grassland. The main crops grown near Agogo are cocoa, coffee, oil palm, citrus, cashew, mango, cassava, yam, among others. The main crops in Wenchi are corn, rice, sorghum, yam, plantain, groundnuts, and cowpea, among others. In Kenieroba, farmers grow mainly rice, sorghum, millet, corn, beans, among other crops; cereals are grown during the wet season (May-October) and vegetables in smaller plots during dry season (November-April).

### Aerial and ground-level insect collection and specimen processing

The aerial collection methods were described in detail previously (Florio et al. 2020; Huestis et al. 2019), Briefly, insect sampling was conducted using sticky nets (panels, each 1 m wide x 3 m long) attached to the tethering line of 3 m diameter helium-filled balloons. Each balloon typically carried three panels, set at 120 m, 160 m, and 190 m agl (above ground level). Balloons were launched around 17:30 before sunset and retrieved around 07:00, after sunrise the following morning. To control for insects trapped near the ground as the panels were raised and lowered, comparable control panels were raised up to 100 m agl and immediately retrieved during each balloon launch and retrieval operation. Following panel retrieval, inspection for insects was conducted in a dedicated clean area. Individual insects were removed from the nets with forceps, counted, and stored in vials containing 80% ethanol. Concurrent samples were collected from a 1 m^2^ panel mounted 1 m above ground to represent ground insects. This net was attached to a line from a frame in such a way that allowed it to orient perpendicular to the wind direction like the nets suspended from the helium balloon.

### Taxon identification

Using a dissecting microscope, African specimens were identified morphologically to order and to morphospecies, counted, and recorded. Specimens of the selected morphotypes were identified by expert taxonomist who narrowed the identification down to species or genus. All African specimens of *C. insularis* were confirmed by Dr. James Zahniser (USDA-APHIS, National Museum of Natural History [USNM], Smithsonian Institution, Washington, DC, USA). Voucher specimens are deposited at USNM.

### Distribution data in the USA

Publicly available records of observations of *C. insularis* from BugGuide.net (n=22; (BugGuide 2024)), iNaturalist (n=827; (iNaturalist 2024)) that met “Research grade” standard (verified by SVH) were downloaded and added to records available from the Florida Department of Agriculture and Consumer Services, Division of Plant Industry (FDACS-DPI) (N=266) after the specimens were identified by FDACS-DPI hemipterists.

Vouchers for FDACS-DPI specimens are deposited in the Florida State Collection of Arthropods. The data from Florida also included 3 interception records in trucks that originated from Arizona (2) and Mexico (1). Border interception records of *C. insularis* (since 2019) from ports of entry in the USA (identified by Dr. James Zahniser, USDA) were provided from USDA/APHIS (N=9) included the port of entry and date of collection. Of 1,124 observations collected until December 31, 2023, from the USA, 11 represented insects during transit, and thus were excluded from distribution data that were used to plot *C. insularis* range in the USA. This compilation of data is included in Table S1.

### Data analysis

Subsamples of the Mali collections from March to December 2019 and the Ghana collections from May to October 2021 were evaluated for the presence of *C. insularis*. Every month of collection at each study site was represented by at least 6 panels at altitude and at least 4 panels at ground level. The total number of insects per panel represents the ‘panel density’. Aerial density was estimated as the panel density of the species divided by the total air volume that passed through that net that night (i.e., aerial density = panel density/volume of air sampled, and volume of air sampled = panel surface area × mean nightly wind speed × sampling duration). The panel surface area was 3 m^2^. Wind-speed data were obtained from the atmospheric re-analyses of the global climate (ERA5). Hourly data consistent of the eastward and northward components (horizontal vectors) of the wind were available at 31-km surface resolution at 2 and 300 m agl (1000 and 975 mbar pressure levels). Overnight records (19:00 through to 06:00) were averaged to calculate the nightly mean direction and mean wind speed over each African sampling station and select locations in the USA (below) based on standard formulae using code written in SAS (SAS software 2019).

The intensity of migration was expressed as the expected number of migrants crossing a line of 1 km perpendicular to the wind direction at altitude, which reflect their direction of movement (Drake, Reynolds 2012; Florio et al. 2020; Hu et al. 2016; Reynolds et al. 2017). We used the mean wind speed at altitude during the migration season (Table 1) and assumed that the leafhoppers fly in a layer depth of 200 m above ground level (Florio et al. 2020). The nightly migration intensity was computed across the flight season (including sampling nights during which no migrants were captured). The corresponding annual index was estimated by multiplying the nightly index by the period of migration, estimated from the first and last month that the species was captured.

**Table 1.** Mean panel and aerial densities of *C. insularis* at altitude and ground levels across study sites with average temperature, humidity, wind speed, and direction during the migration period (May-August).

|  | Agogo (panels: 66/19)<sup>a</sup> |  | Wenchi (panels: 57/26) |  | Kenieroba (panels: 47/0) |  |
| --- | --- | --- | --- | --- | --- | --- |
|  | Altitude (26)<sup>b</sup> | Ground (21)<sup>b</sup> | Altitude (19) | Ground (12) | Altitude (35) | Ground (12) |
| Panel density *C. insularis*<sup>c</sup> | 0.23 (0-0.47) | 0.05 (0-0.15) | 0.40 (0.0-0.84) | 0.17 (0-0.53) | 0 | 0 |
| Total *C. insularis* /total insects<sup>d</sup> | 0.26 (6/2314) | 0.06 (1/1662) | 0.21 (8/3852) | 0.17 (2/1193) | 0 (0/3348) | 0% (0/549) |
| Aerial density *C. insularis*<sup>e</sup> | 0.41 (0-0.83) | nd | 1.0 (0-2.08) | nd | 0 | 0 |
| Dispersal mass (n/[km night])<sup>f</sup> | 17,004 (0-34,422) | nd | 35,424 (0-73,682) | nd | 0 | 0 |
| Wind speed (m/s)<sup>g</sup> | 4.75 (4.6-4.9) | 1.86 (1.8-1.9) | 4.07 (3.96-4.18) | 1.85 (1.8-1.9) | 1.91 (1.8-2.0) | 1.91 (1.8-2.0) |
| Wind direction (°)<sup>g</sup> | 244 (243.7-244.2) | 246.3 246.1-246.6) | 244.2 (244.0-244.5) | 246.8 (246.5-247.2) | 229.2 (229.0-229.4) | 229.1 (229.0-229.4) |
| Temperature (°C)<sup>g</sup> | 23.9 (23.8-24.1)[19.3] | 25.3 (25.1-25.4)[20.7] | 24.9 (24.7-25.1)[19.9] | 26.4 (26.2-26.5)[21.3] | 28.0 (27.6-28.4)[17.6] | 29.5 (29.1-29.9)[19.5] |
| Relative humidity (%)<sup>g</sup> | 85.0 (84.1-85.8) | 84.1 (83.2-84.9) | 80.2 (79.2-81.2) | 80.8 (79.8-81.8) | 75.6 (73.5-77.7) | 75.6 (73.5-77.7) |
| <sup>a</sup> Total number of panels/control panels inspected |  |  |  |  |  |  |
| <sup>b</sup> Number of panels |  |  |  |  |  |  |
| <sup>c</sup> Mean (95%CI) panel density of *C. insularis* /panelr during May-September (zeros replace negative lower 95%CL). |  |  |  |  |  |  |
| <sup>d</sup> Percentage of *C. insularis* from total insects (% total *C. insularis* /total insects) |  |  |  |  |  |  |
| <sup>e</sup> Mean (95%CI) aerial density of *C. insularis* /10<sup>6</sup> m<sup>3</sup> of air during May-September (zeros replace negative lower 95%CL). |  |  |  |  |  |  |
| <sup>f</sup> Estimated number of *C. insularis* crossing an imaginary 1 km line perpendicular to the wind direction between 50-250 m above ground level |  |  |  |  |  |  |
| <sup>g</sup> Nightly hourly averages and 95%CL (over May-August; N = 2,706) [minimum temp] |  |  |  |  |  |  |

To assess the likelihood of windborne movement to a new locations in the USA, we identified new sites where *C. insularis* was observed for the first time outside its previous year’s range, defined by connecting all the extreme points of its cumulative distribution in the previous year. For each new site, we consider its nearest known site—where *C. insularis* was previously reported—as a putative source. Underlying our approach is the assumption that the missing data due to low sampling in certain localities and/or certain years would generate noisier data-patterns rather than as systematic pattern. Therefore, finding a consistent biological trend (signal) in these data, relevant to the process in question is likely produced by a biological process rather than by variation in sampling intensity. Virtually all datasets on geographic expansion at these scales of time and space would present similar “imperfections”, inviting inquiries to better assess and address their limitations. However, large Citizen-Scientist databases provide compelling advantages as pointed out by (Kittelberger et al. 2021).

The annual distributions of nightly (19:00—06:00) winds during the year of the new record were plotted as vectors pointing to the direction the insects would be carried if they flew 8 hours from that site on that night’s wind at 300 m agl. The self-propelled flight speed of leafhoppers does not typically exceed 1 m s^−1^ (Zhou et al. 2003) and therefore at altitude with winds exceeding 4 m/s, the direction of the movement of the insects will be determined primarily by the wind. Only nights with at least 8 hours when temperatures were >16^○^C at altitude and on the ground were included (assuming no flight occurred below 16^○^C (Shields, Testa 1999). The total flight duration of tethered leafhoppers reached over 7 h (Zhou et al. 2003) and additionally, inferences about windborne leafhoppers in the US traveling hundreds of kilometers per night (or tens of kilometers/night over successive nights) have been made based on their phenology (Carlson et al. 1992; Reynolds et al. 2017; Shields, Testa 1999). Thus, we assumed that *C. insularis* could fly at altitude between 2 and 8 h per night. The wind trajectories were projected on maps based on the wind direction, speed, and 8 hours of flight using SAS 9.4 (SAS software 2019). Windborne dispersal from the putative source to its destinations was assessed based on visual inspection of the trajectory distributions.

## Results & Discussion

### African aerial and ground surveillance

Overall, 25,431 insect specimens collected in West Africa on 308 panels (157 panels at 120-290 m agl, 84 panels at 1 m agl, and 67 control panels) were sorted and evaluated for the presence of *Curtara insularis*. Interception of *C. insularis* at altitude and at ground level occurred between May and August (Fig. 1) but not in September and October (sampling in other months were only performed in Mali). Although we cannot rule out high-altitude flights in the rest of the year, we consider the period between May and August as its migration period and unless otherwise stated, confined the summary statistics to this period (Table 1, Fig. 1). A total of 18 specimens of *C. insularis* were identified among samples collected in Agogo and Wenchi (Ghana) during May to August (9,113 insect specimens, Table 1), but none was found in Kenieroba (Mali) during these months (3,897 insect specimens, Table 1) or throughout the year (6,151 insect specimens). Most *C. insularis* (N=14) were intercepted at altitude (160-190 m agl), 3 were collected at ground level, and 1 on a control panel (Table 1). A single specimen on the control panels (N=45 control panels) as opposed to 14 specimens intercepted in these locations at altitude (N=61 panels, Table 1) indicate that “contamination” near the ground (<50 m agl) is unlikely to account for the capture of so many specimens at altitude. At altitude, similar numbers of males (8) and females (6) were collected, indicating that both sexes equally engage in high-altitude flight unlike mosquitoes, in which females consist of >80% from the aerial collection (Huestis et al. 2019; Yaro et al. 2022). All specimens were macropterous (having fully formed fore- and hindwings), albeit no polymorphism in wing development has been noted for this species.

The higher numbers of *C. insularis* at altitude and its relative larger fraction of the total insects on the panels among all specimens collected (Table 1) attest for its propensity to engage in high-altitude windborne dispersal. The scale of *C. insularis* movement at altitude was estimated as the average number of individuals flying between 50 and 250 m agl crossing a 1 km line perpendicular to the wind during a single night. With tens of thousands flying nightly at altitude across 1 km (Table 1), the expected number over the four-months season (Fig. 1) was in the millions, reflecting that this species had established robust populations in its source sites. Radar studies have shown that windborne insects fly nightly over swathes that are tens or even several hundreds of kilometers wide (Drake, Reynolds 2012; Hu et al. 2016). With a mean wind speed of 4.5 m/s, individual insects flying 2-8 hours would readily cover 30-150 km per night. High-altitude windborne dispersal has been reported in leafhoppers including a flight >3,000 km over the ocean (*Balclutha salturella* [Previously: *B. pauxilla*]), culminating in the invasion of Ascension Island (Ghauri 1983), and seasonal migrations over hundreds of kilometers in North America (*Empoasca fabae, Macrosteles quadrilineatus*, and *Circulifer tenellus*), and Asia (*Nilparvata lugens* and *Sogatella furcifera*) (Carlson et al. 1992; Glick 1939; Hu et al. 2019; Reynolds et al. 2017; Taylor 1974; Wu et al. 2019). In tropical West-Africa, temperature and humidity rarely restrict flight activity even at altitude (Table 1; (Sanogo et al. 2021)). Altogether, high-altitude dispersal of *C. insularis* is a potent strategy for spreading rapidly over large areas. These large number of migrants account for massive propagule pressure (Lockwood et al. 2005; Simberloff 2009) that could well explain the importance of this modality of dispersal compared with vehicular spread, which involves small number of specimens that often arrive in inhospitable spaces, e.g., storage facilities surrounded by urban environment, which limit reproduction success and establishment of populations.

Given the predominant wind directions in this region of West Africa (Fig. 1, Table 1), *C. insularis* is probably carried by the winds inland, encroaching up to at least ∼300 km (Wenchi) from its supposed landing sites in one or more West Africa ports (Fig 1). If it had been introduced through the largest port of West Africa, Abidjan, Ivory Coast, the predominant southwesterly winds would have carried it straight to the Agogo and Wenchi areas (Fig. 1), whereas Kenieroba is clearly off the predominant wind trajectories from the main Gulf of Guinea ports. Thus, invasion of locations such as Kenieroba may require over-land vehicular transport or considerably longer time using windborne dispersal. Whether *C. insularis* can establish populations in Kenieroba, Mali is an open question. Located within the Sudano-Guinean zone along the Niger river, the overall climatic conditions in Kenieroba are similar to those in Wenchi (Table 1) or other sites where *C. insularis* is found (below), albeit with a longer dry season (December-April). Other explanations in which *C. insularis* has arrived earlier at other region(s) of Africa cannot be rule out without additional data on its distribution on the continent.

### Spread of *Curtara insularis* in the USA

The current distribution of *C. insularis* in the USA is based on 1,109 records (excluding 11 “in-transit” records) spanning the period of 2004-2023 (Fig. S2 and Table S1). The records cover much of the southeastern and south-central USA (25°—36°N latitudes and 77°— 102°W longitude, Fig. S2). There was minimal seasonal variation considering the month of observation and latitude or longitude (not shown), suggesting that *C. insularis* populations are perennial in that range. Given the number of the records, this range suggests a habitat-suitability space, which appears bounded to the north by annual temperature minima isotherm of −12°C, described by vegetation zone 8 (USDA 2023) and to the west by annual precipitation isohyet of 510-630 mm (NOAA 2023). Despite their apparent correlation between these boundaries and the current limits on the *C. insularis* distribution, studies are needed to establish causal relationships of these hypothetical factors. Permissive combinations of temperatures and precipitation also occur along the Northwest coast (e.g., in California) and the Southwest (e.g., in Arizona). Indeed, *C. insularis* was intercepted in Florida on two trucks that originated from Arizona in 2022 and 2024 (FDACS-DPI records), although the species has not yet been reported from any western state. Future records of this species from those territories may reveal the stability of the current range. We consider the range of records available up to December 2023 as the space in which range expansion could have occurred, evaluating how *C. insularis* “filled” it.

The interception of *C. insularis* in international ports (9 records since 2019) and on trucks entering Florida (3 records since 2004) provide evidence for the role of the maritime trade as well as vehicles overland in transporting this species. The port interceptions were in Florida (5), Houston Texas (2), Georgia (1), and Puerto Rico (1). These records substantiate that *C. insularis* can spread by all these means as well as by wind at altitude (above) as other invasive pests, such as *Helicoverpa* and *Spodoptera* moths (Jones et al. 2019). Given the large distances of transport by truck (daily average of ∼1,000 km), airplane, or ships, the expected pattern of spread would be by long and irregular leaps in all directions (Ahn et al. 2023; Lehmann et al. 2023; Suarez et al. 2001), whereas windborne spread would predict a continuous incremental spread that resembles “diffusion process” that follows predominant winds.

The first record of *C. insularis* in the USA dates to January 2004 (Hillsborough County, Florida, Table S1). However, 42 specimens from 5 central-south Florida counties were identified that year, indicating the original introduction took place into that region earlier (Fig. 2A Fig. S3). Over the next 5 years, it has spread throughout the state, reaching the panhandle by 2008, and expanding outside the state only in 2012 (Fig. S3). The relatively slow spread northward, given the numerous sightings in Florida and compared with subsequent years is incompatible with vehicular transport but can be attributed to “unfavorable” predominant winds which have a strong east-west component in relation to the narrow peninsula (∼200 km wide), resulting in most windborne journeys ending at sea (Fig. 2A). That Florida is a major producer of most nursery stocks and diverse produce that are shipped by trucks efficiently all over the US, highlights the contrast between this mode of transport and the pattern of spread observed. In 2012, *C. insularis* was reported near Houston, Texas, and a year later near Lafayette, Louisiana (Fig. S3). With average nightly wind speeds 5.6 m/s (maximum of 16.9 m/s) we expect displacement ranges of 40-150 km/night over 2-8 hours of flight with up to 500 km during maximum recorded (nightly) windspeed. The closest *C. insularis* record was from Florida, >900 km away from Houston (500 km away from Lafayette). The scarcity of westward winds from Florida and the large distance suggests that *C. insularis* was transported to Houston by a ship, truck, or airplane rather than by the wind (Fig. 2B, and below). Nonetheless, its arrival near Lafayette (Louisiana, Fig. S3) the following year is consistent with frequent eastward winds from Houston (Fig. 2B). In 2016, *C. insularis* was observed around Austin and Dallas (Texas) as well as near Columbus, Georgia (Fig. S3). Wind trajectories from Houston to Dallas were common and those towards Austin are modestly common, as are the trajectories from Tallassee to Columbus (Fig. 2B). The slower arrival into these destinations (∼4 years) agrees with the modest frequency of corresponding wind trajectories. By 2017 and 2018, *C. insularis* was reported from South Carolina and in new counties of Texas, including a remote region of western Texas >50 km from a major highway (Fig. S3). Fifteen of the new records were located consistent with wind trajectories from previously established populations (Fig. 2C), however 4 of 20 would require 9-11 hours windborne flight (at average nightly wind speed) and a single site near Matamoros, Mexico, would require ∼16 hours flight or an additional landing by ship at the nearby port of Brownsville Texas (Fig. 2C, S3). In 2019, *C. insularis* was reported from its northern-most point near Raleigh, North Carolina (∼170 km from Florence, SC Fig. S3), Tuscaloosa, Alabama (∼260 km from Crestview FL), and Jackson, Mississippi (∼130 km from Pensacola FL), as well as to other new localities, closer to previously established populations (Fig. 2d, S3). All these sites were consistent with common wind directions and distances from previously established populations (Fig. 2d, S3). Similarly, new counties in 2020 and 2021-2023 were also <100 km from previously established sites and often from multiple such sites (Fig. 2d S3), consistent with windborne spread. This approach is conservative because it assumes that the source populations of news localities are known, although unknown source sites may be closer to the new location, resulting in overestimating the distance these migrants actually passed. The average annual maximum distance of range expansion was 229 km (range: 0-950 km, Fig. S4). The arrival into Houston (recorded on 2012) presented an extreme outlier, indicating a different mode of range expansion from the rest (Figs. 2, S3, and S4). Excluding this value, yields an average of annual maximum range expansion of 176 km (Fig. S4: inset). Both values appear considerably lower than the average maximum distances of truck transporting produce in the US.

**Figure 2.**
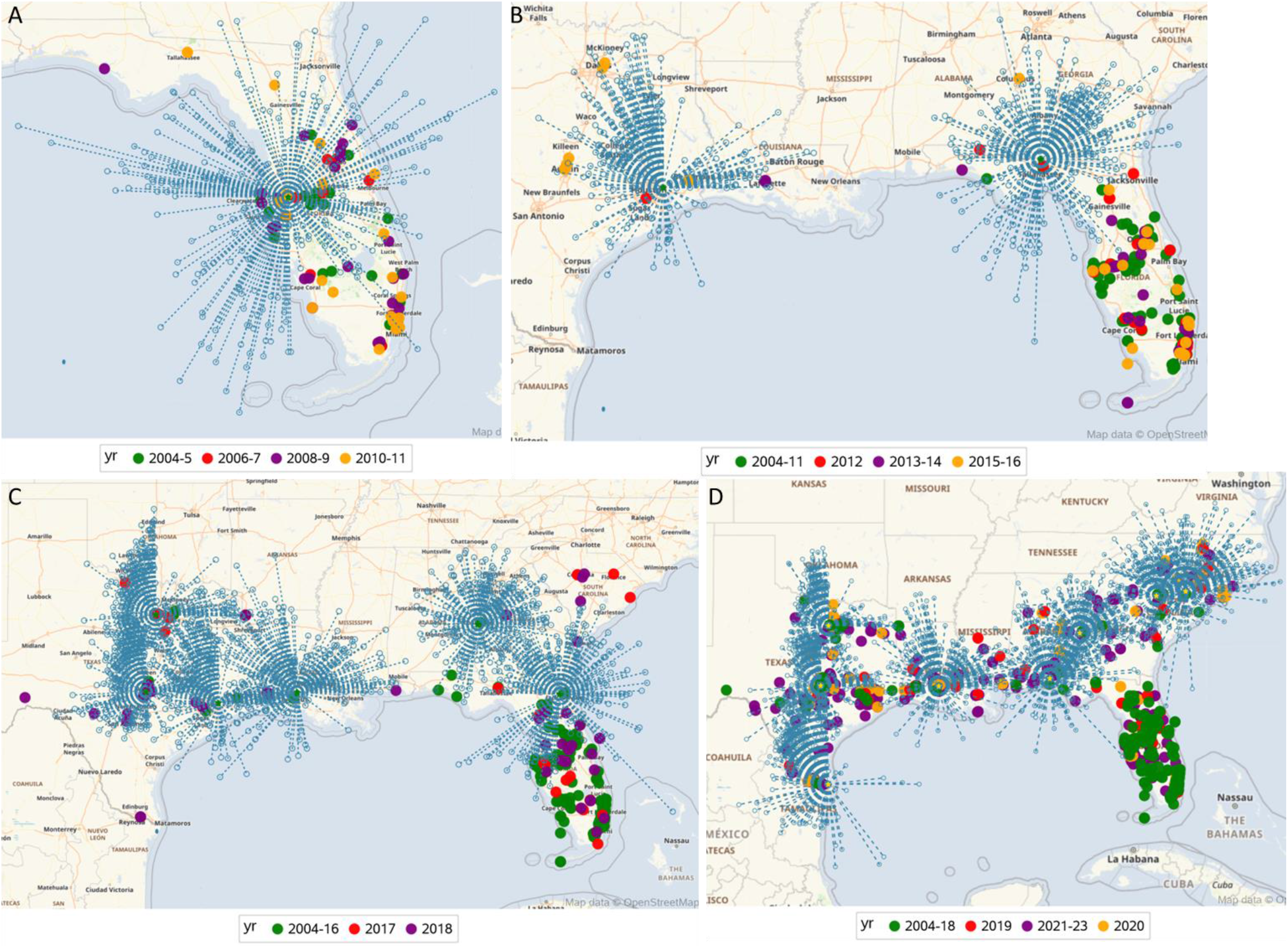
Maps showing range expansion of *Curtara insularis* in relation to projection of annual wind trajectories at altitude (300 m asl) from putative source(s) reported previously assuming 8-hour windborne flight and linear winds (broken lines). A) Expansion of *C. insularis* (2004— 2011) with annual nightly projections of wind trajectories from Tampa 2005 (see text). B) Expansion of *C. insularis* (2012—2016) with annual nightly wind trajectories from Tallahassee and Houston 2012 (see text). C) Expansion of *C. insularis* (2017—2018) with projections of annual nightly wind trajectories from Jacksonville, Tallahassee, Columbus, Lafayette, Houston, Dallas, and Austin. Expansion of *C. insularis* (2018—2023) with projections of annual nightly wind trajectories from Florence, Columbia, Columbus, Crestview, Lafayette, Dallas, Austin, Matamoros (Mexico).

### Conclusion and implications

Our results show that *C. insularis* exploits multiple modes of long-range dispersal, including vehicular transportation on board of ships and trucks and windborne migration at altitude. *Curtara insularis* was first found in Africa by sampling at altitude. Based on its aerial density over Ghana, we estimate that annually, millions of *C. insularis* migrate at altitude across each 1 km sections perpendicular to the wind, representing a massive propagule pressure that probably exceeds by several orders of magnitude that of transport by vehicles. Extending these findings, we assess the relative importance of windborne migration compared with vehicular transport to the spread of *C. insularis* in the USA. Given the size of the habitat space this leafhopper has expanded to until 2023, it is notable that it took 5 years to reach the Florida panhandle from central Florida (∼350 km), and 8 years to spread beyond Florida to other states. Likewise, all but three of the hundreds of records of inter-annual range-expansion exceed 300 km from the nearest previously known site. Had it regularly been transported by trucks (or airplanes) overland to the numerous new areas it was reported from, it would have spread from Central Florida and reach North Carolina (∼1,000 km), or western Texas (∼2,000 km) considerably faster than 19 or 15 years, respectively. Our data suggest that except 1-3 independent introductions to the US by the maritime trade (and possibly by trucks), *C. insularis* expansion overland has been incremental, diffusion-like process, which generally agrees with common wind trajectories. Thus, *C. insularis* range expansion in the US is better explained by high-altitude windborne dispersal following one or few successful journeys onboard ships. How unique is *C. insularis* among invasive insects in exploiting high-altitude windborne dispersal? Because strong dispersive capacity is a key trait of invasive species (Anderson et al. 2010; Hu et al. 2019; Hulme et al. 2008; Jones et al. 2019; Lounibos 2002; Mack et al. 2000; Renault et al. 2018) and because of the diverse insect faunas at altitude (Chapman et al. 2004; Drake, Reynolds 2012; Florio et al. 2020; Hu et al. 2016; Huang et al. 2024; Yaro et al. 2022), we propose that-high altitude windborne dispersal may well be especially common among invasive insect species. Some of these species pose severe risks to biodiversity, food security, and public health (above), and are exacerbated by anthropogenic changes including climate change (Cao, Feng 2024). Therefore, aerial surveillance 10-30 km downwind from major ports might complement traditional surveillance procedures to discover invading insect species early and predict their likely destinations—information that can improve elimination efforts before they spread over large areas—when elimination may be an especially viable option.

## Acknowledgements

We thank Drs. James Zahniser for identification of the *Curtara insularis* specimens from our collection and for discussions on this manuscript, Adilson Pinedo-Escatel for helpful comments on this manuscript, Carolina Barillas-Mury, Jesus Valenzuela, Thomas Wellems, Mrs. Fatoumata Bathily, and Mrs. Margery Sullivan for their support. Special thanks to the residents of the villages Kenieroba, Dukusen, and Wenchi for their cooperation and hospitality. This study was supported by the Division of Intramural Research, National Institute of Allergy and Infectious Diseases, National Institutes of Health, Bethesda MD (ZIA AI001196-06), the Bill & Melinda Gates Foundation, Grand Challenges Explorations grant (OPP1217659) awarded to TL, the Florida Department of Agriculture and Consumer Services, the United Kingdom Biotechnology and Biological Sciences Research Council to Rothamsted Research, Division of Plant Industry, and the U.S. Department of Agriculture, Agricultural Research Service. Mention of trade names or commercial products in this publication is solely for the purpose of providing specific information and does not imply recommendation or endorsement by the USDA. USDA is an equal opportunity provider and employer.

## Supplementary materials

Three observations of *Curtara* spp. in coastal Ghana made between August and December 2021 (iNaturalist observations 92716825, 103237938, 104023032), preceded by an observation in coastal Benin in December 2020 (iNaturalist observation 65165749), and one near Lagos, Coastal Nigeria in 2022 (iNaturalist observation 113460502), suggest multiple ports of entry in Africa. These records likely represent additional records of *Curtara insularis* (Caldwell), however, specimens will need be identified with certainty, as there are externally similar species in the *Curtara samera*-group as defined by DeLong and Freytag (1976) and the scarcity of these records.

**Fig. S1.**
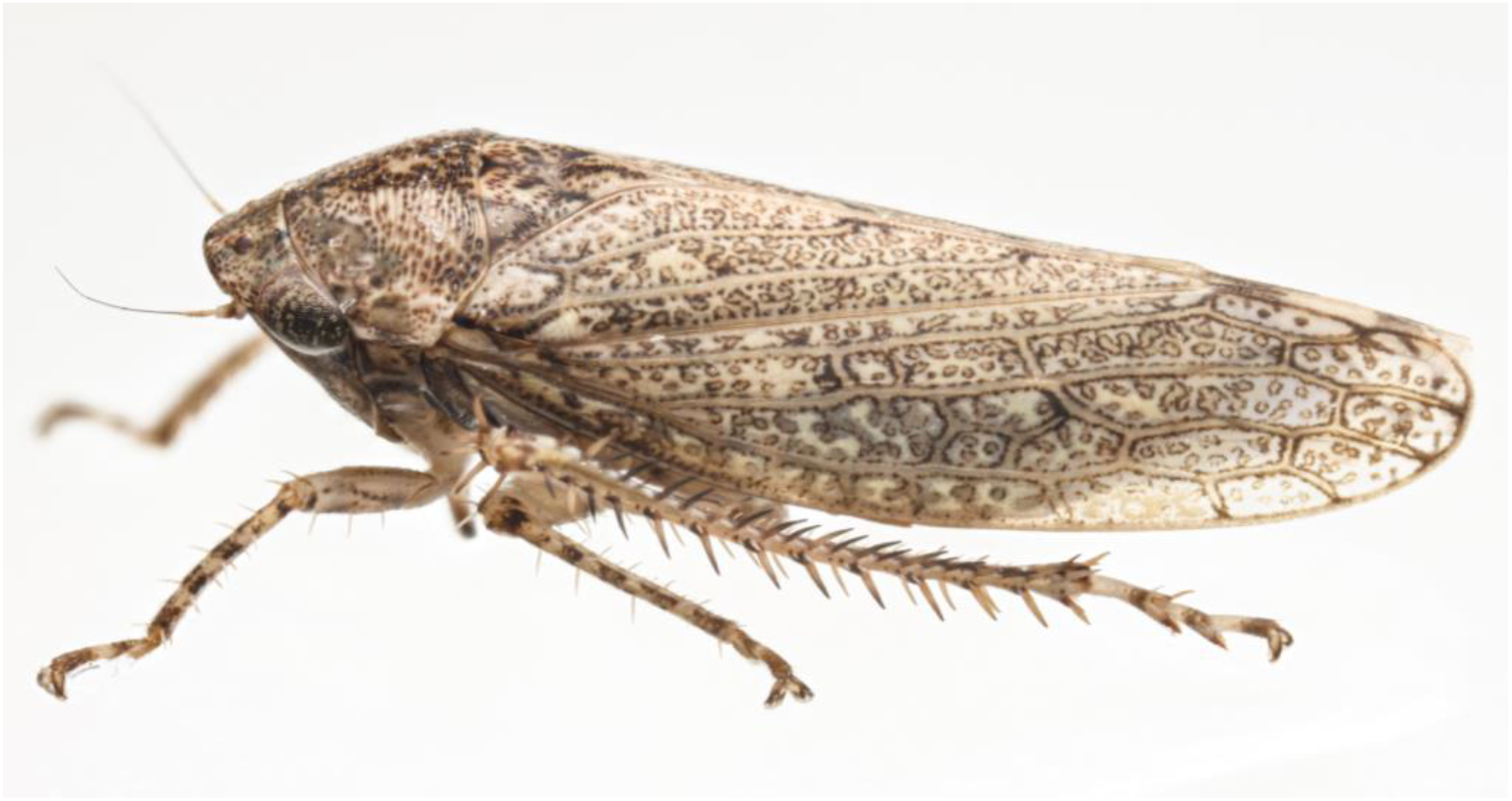
*Curtara insularis*: lateral view showing reddish ring spots on wings (source: Solomon V. Hendrix)

**Fig. S2.**
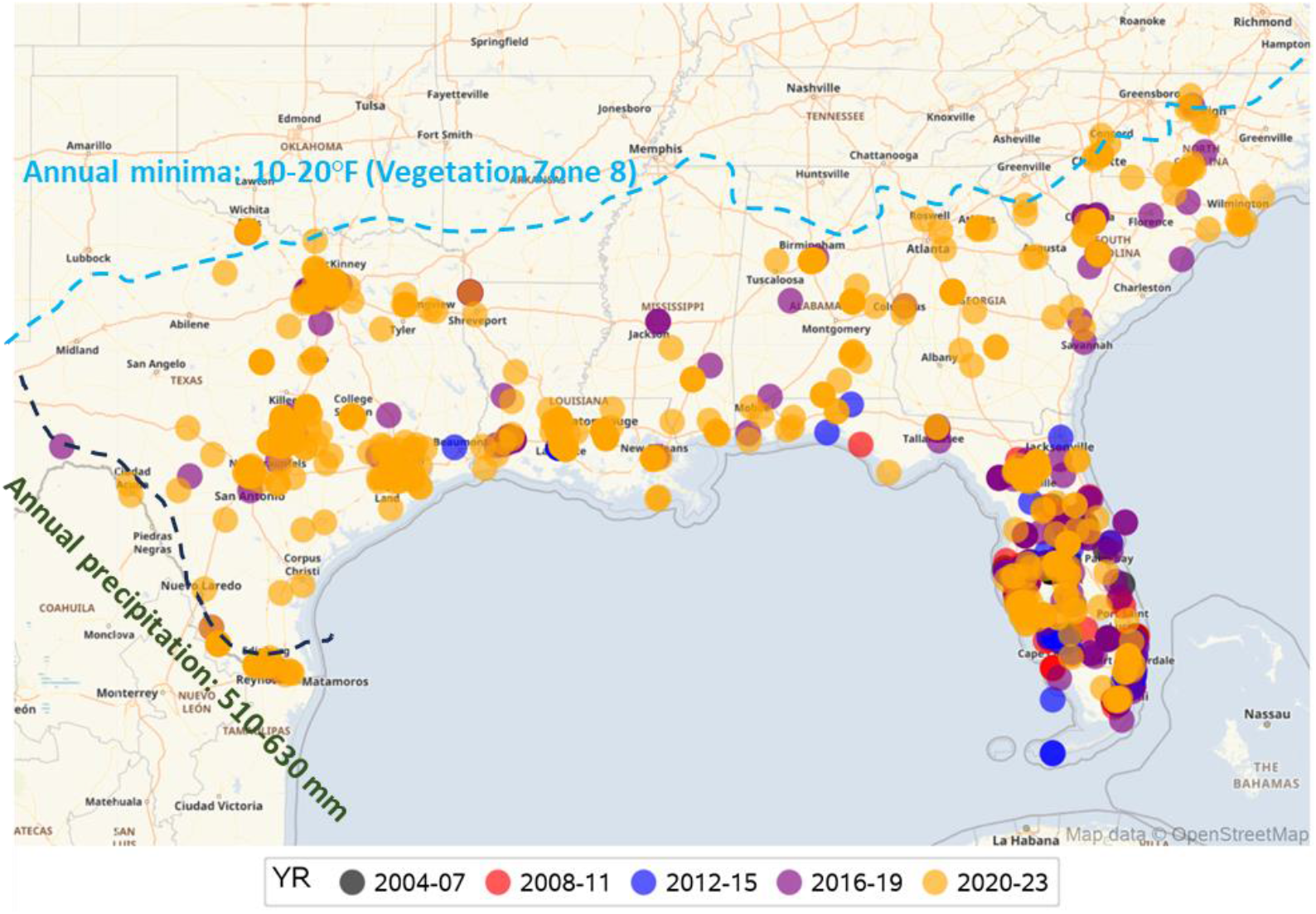
Map showing distribution of *C. insularis* in the USA based on the records compiled in this paper (Table S1) by four-year intervals. Blue line is schematically drawn to represent annual temperature minima isotherm of −12°C (USDA vegetation zone 8 (USDA 2023) and the black line was drawn to represent the annual precipitation isohyet of 510-630 mm (NOAA 2023).

**Figure S3:**
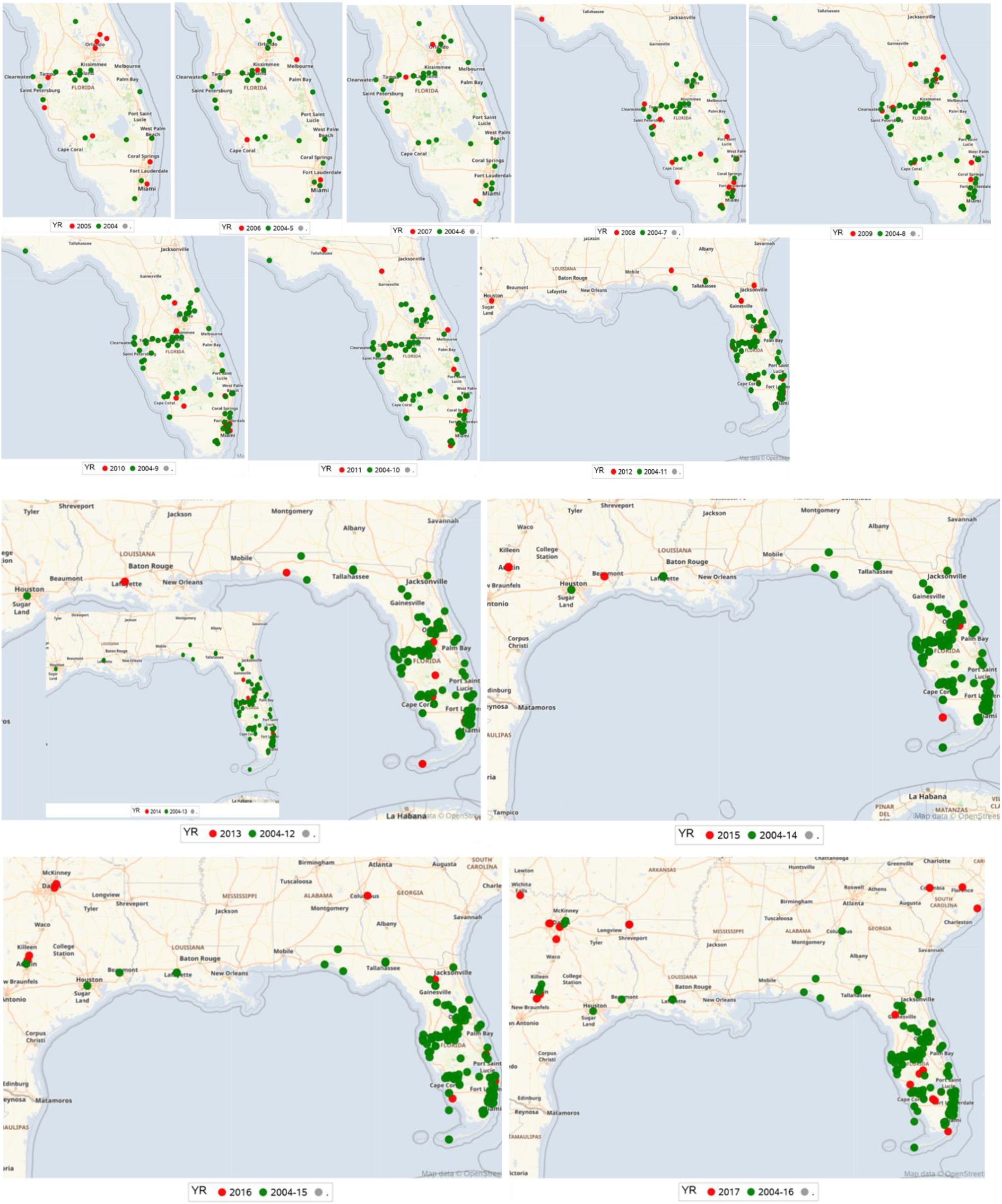
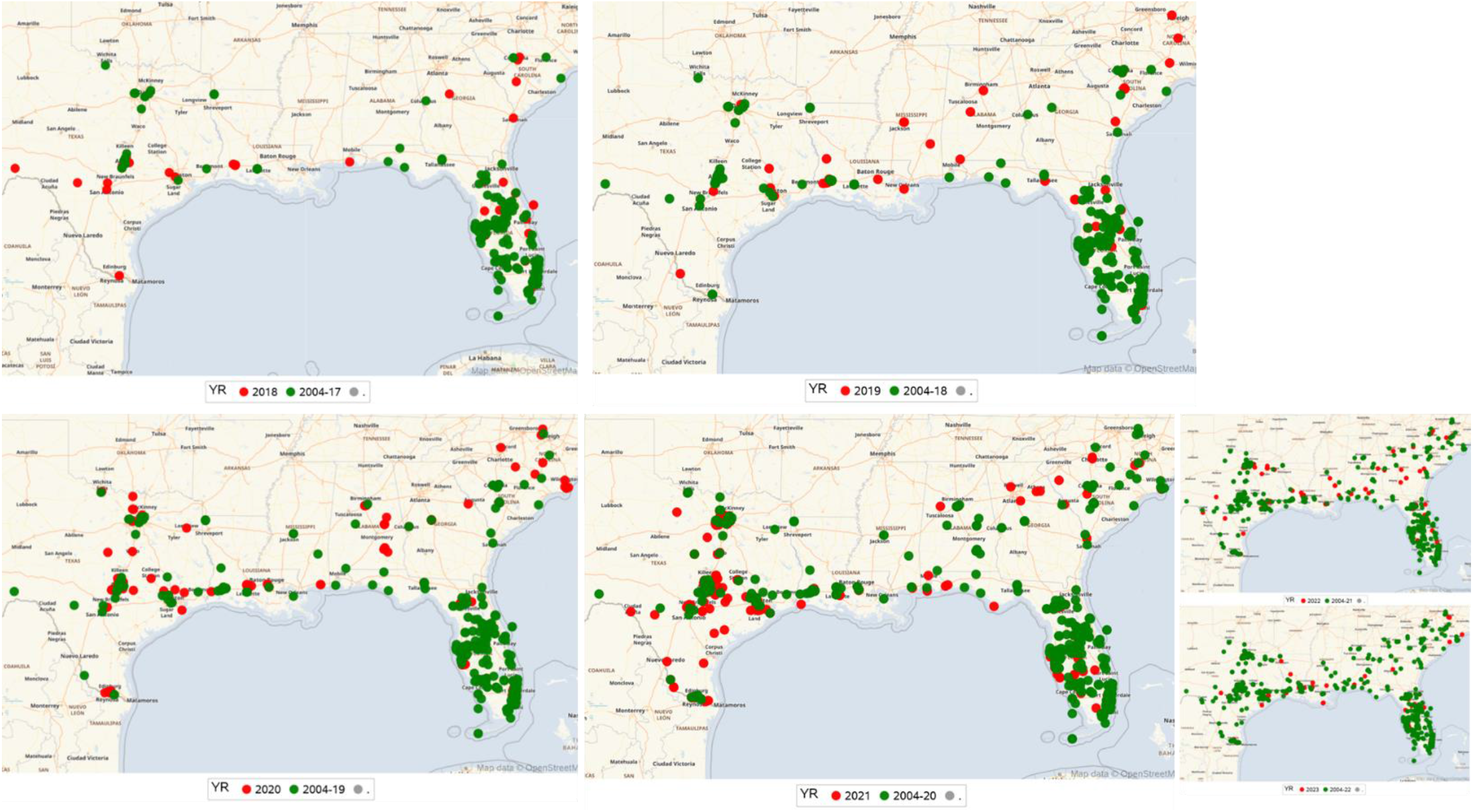
Annual range expansion of *C. insularis* showing each year new records (red) and cumulative previous records (green). Range expansion is defined as location outside the polygon connecting all extreme sites (Methods). Map dimension changes based on sighting locations and years with minimal range expansion are shown as inset (2014) or reduced in size (2022-3).

**Figure S4:**
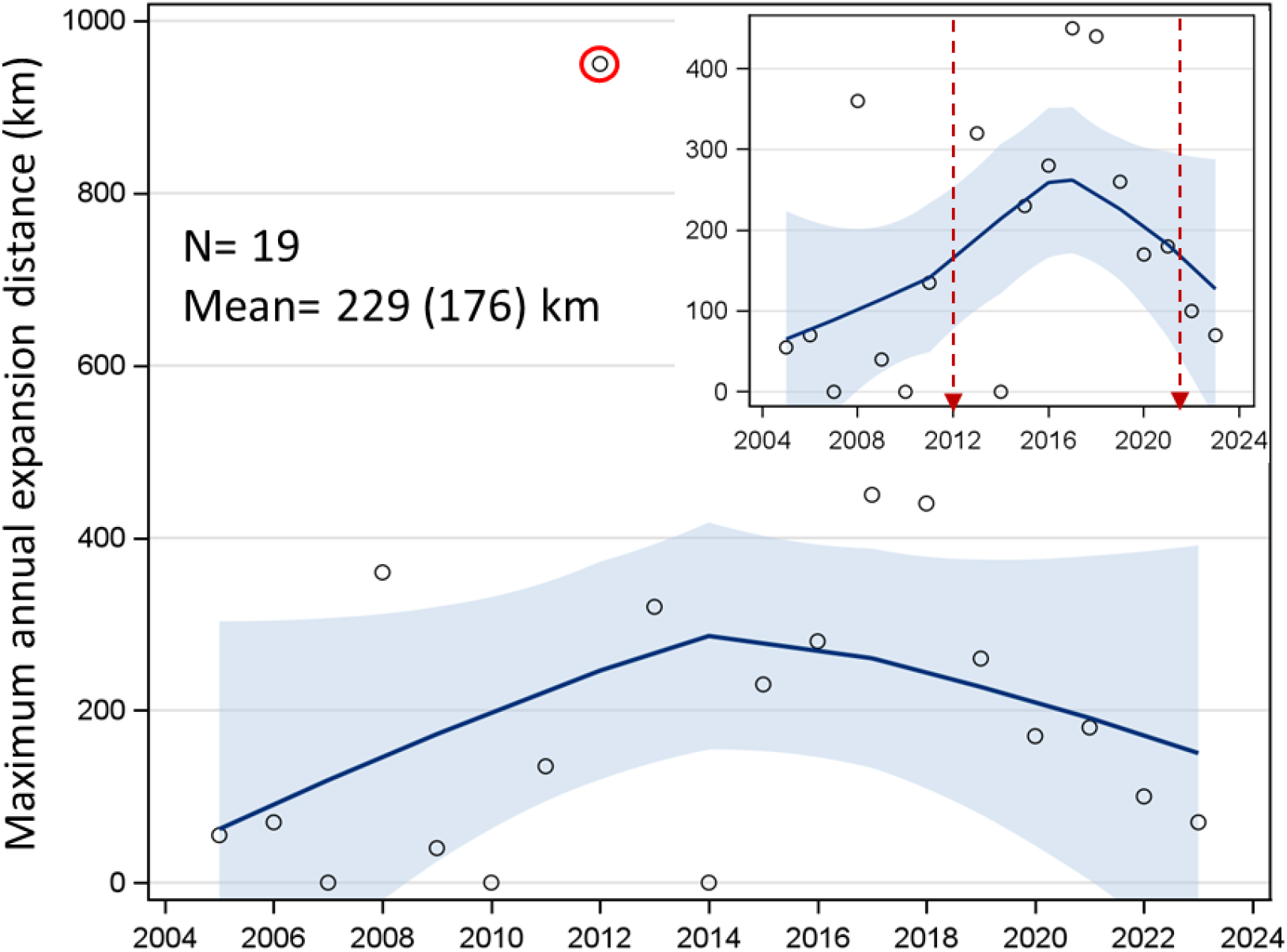
Over time variation in the maximum annual range-expansion distance measured between each year new sightings and the closest previous sightings. Red circle around the extreme outlier (2012) shows a distance of 950 km from the nearest previous record. Inset: The same graph excluding the 2012 extreme outlier. Arrows separate periods of apparent larger annual expansion distances.

## References

Ahn J, Sinka M, Irish S, et al. (2023) Modeling marine cargo traffic to identify countries in Africa with greatest risk of invasion by Anopheles stephensi. Sci Rep 13:876

Almeida RPP, Nunney L (2015) How Do Plant Diseases Caused by Xylella fastidiosa Emerge? Plant Disease 99:1457–1467

Anderson KL, Deveson TE, Sallam N, et al. (2010) Wind-assisted migration potential of the island sugarcane planthopper (Hemiptera: Delphacidae): implications for managing incursions across an Australian quarantine frontline. J Appl Ecol 47:1310–1319

BugGuide (2024). Iowa State University,

Cao RY, Feng JM (2024) Future Climate Change and Anthropogenic Disturbance Promote the Invasions of the World’s Worst Invasive Insect Pests. Insects 15

Carlson JD, Whalon ME, Landis DA, et al. (1992) Springtime Weather Patterns Coincident with Long-Distance Migration of Potato Leafhopper into Michigan. Agr Forest Meteorol 59:183–206

Chapman JW, Reynolds DR, Smith AD, et al. (2004) An aerial netting study of insects migrating at high altitude over England. B Entomol Res 94:123–136

Chapman JW, Reynolds DR, Wilson K (2015) Long-range seasonal migration in insects: mechanisms, evolutionary drivers and ecological consequences. Ecology Letters 18:287–302

Drake VA, Reynolds DR (2012) Radar Entomology: Observing Insect Flight and Migration. CABI International

Elton CS (1958) The ecology of invasions by animals and plants. Chapman and Hall, London

Florio J, Verú LM, Dao A, et al. (2020) Diversity, dynamics, direction, and magnitude of high-altitude migrating insects in the Sahel. Sci Rep-Uk 10

Fu XW, Li C, Feng HQ, et al. (2014) Seasonal migration of Cnaphalocrocis medinalis (Lepidoptera: Crambidae) over the Bohai Sea in northern China. Bull Entomol Res 104:601–9

Ghauri MSK (1983) A case of long-distance dispersal of a leafhopper. In: Robertson TS, Wilson MR, Knight WJ and Pant NC (eds) 1st International Workshop on Leafhoppers and Planthoppers of Economic Importance. Commonwealth Institute of Entomology, London, London, UK, pp. 249–255

Glick PA (1939) The Distribution of Insects, Spiders and Mites in the Air Technical Bulletin. United States Department of Agriculture, Washington, D.C., 151 pp., pp. 151

Hu G, Lim KS, Horvitz N, et al. (2016) Mass seasonal bioflows of high-flying insect migrants. Science 354:1584–1587

Hu G, Lu MH, Reynolds DR, et al. (2019) Long-term seasonal forecasting of a major migrant insect pest: the brown planthopper in the Lower Yangtze River Valley. J Pest Sci (2004) 92:417–428

Huang J, Feng H, Drake VA, et al. (2024) Massive seasonal high-altitude migrations of nocturnal insects above the agricultural plains of East China. Proc Natl Acad Sci U S A 121:e2317646121

Huestis DL, Dao A, Diallo M, et al. (2019) Windborne long-distance migration of malaria mosquitoes in the Sahel. Nature 574:404–408

Hulme PE, Bacher S, Kenis M, et al. (2008) Grasping at the routes of biological invasions:: a framework for integrating pathways into policy. J Appl Ecol 45:403–414

iNaturalist (2024) iNaturalist community.

Jones CM, Parry H, Tay WT, et al. (2019) Movement Ecology of Pest Helicoverpa: Implications for Ongoing Spread. Annu Rev Entomol 64:277–295

Kittelberger KD, Hendrix SV, Sekercioglu CH (2021) The Value of Citizen Science in Increasing Our Knowledge of Under-Sampled Biodiversity: An Overview of Public Documentation of Auchenorrhyncha and the. Front Env Sci-Switz 9

Lehmann T, Bamou R, Chapman JW, et al. (2023) Urban malaria may be spreading via the wind-here’s why that’s important. Proc Natl Acad Sci U S A 120:e2301666120

Li XJ, Wu MF, Ma J, et al. (2020) Prediction of migratory routes of the invasive fall armyworm in eastern China using a trajectory analytical approach. Pest Manag Sci 76:454–463

Lockwood JL, Cassey P, Blackburn T (2005) The role of propagule pressure in explaining species invasions. Trends Ecol Evol 20:223–228

Lounibos LP (2002) Invasions by insect vectors of human disease. Annual Review of Entomology 47:233–266

Mack RN, Occhipinti A (1999) Biotic invasion: A global perspective and ecology of invasion: Patterns and perspectives. Perspectives in Ecology: 67–74

Mack RN, Simberloff D, Lonsdale WM, et al. (2000) Biotic invasions: Causes, epidemiology, global consequences, and control. Ecol Appl 10:689–710

NOAA (2023) U.S. Climate Atlas. In: National Centers for Environmental Information. https://www.ncei.noaa.gov/access/climateatlas/ Accessed: March 21, 2024 2024

Pysek P, Hulme PE, Simberloff D, et al. (2020) Scientists’ warning on invasive alien species. Biol Rev 95:1511–1534

Renault D, Laparie M, McCauley SJ, et al. (2018) Environmental Adaptations, Ecological Filtering, and Dispersal Central to Insect Invasions. Annual Review of Entomology, Vol 63 63:345–368

Reynolds DR, Chapman JW, Stewart AJA (2017) Windborne migration of Auchenorrhyncha (Hemiptera) over Britain. Eur J Entomol 114:554–564

Reynolds DR, Smith AD, Mukhopadhyay S, et al. (1996) Atmospheric transport of mosquitoes in northeast India. Med Vet Entomol 10:185–6

Riley JR, Reynolds DR (1996) Preferential flight by desert locusts on favourable winds. 22nd Conference on Agricultural & Forest Meteorology with Symposium on Fire & Forest Meteorology/12th Conference on Biometeorology & Aerobiology:436–437

Riley JR, Reynolds DR, Smith AD, et al. (1995) Observations of the Autumn Migration of the Rice Leaf-Roller Cnaphalocrocis-Medinalis (Lepidoptera, Pyralidae) and Other Moths in Eastern China. B Entomol Res 85:397–414

Sanogo ZL, Yaro AS, Dao A, et al. (2021) The Effects of High-Altitude Windborne Migration on Survival, Oviposition, and Blood-Feeding of the African Malaria Mosquito, Anopheles gambiae s.l. (Diptera: Culicidae). J Med Entomol 58:343–349

SAS software (2019) SAS Institute Inc. Cary, NC, USA,

Shields EJ, Testa AM (1999) Fall migratory flight initiation of the potato leafhopper, (Homoptera: Cicadellidae): observations in the lower atmosphere using remote piloted vehicles. Agr Forest Meteorol 97:317–330

Siebert A (2014) Hydroclimate Extremes in Africa: Variability, Observations and Modeled Projections. Geogr Compass 8:351–367

Simberloff D (2009) The Role of Propagule Pressure in Biological Invasions. Annu Rev Ecol Evol S 40:81–102

Sinka ME, Pironon S, Massey NC, et al. (2020) A new malaria vector in Africa: Predicting the expansion range of Anopheles stephensi and identifying the urban populations at risk. Proc Natl Acad Sci U S A 117:24900–24908

Suarez AV, Holway DA, Case TJ (2001) Patterns of spread in biological invasions dominated by long-distance jump dispersal: Insights from Argentine ants. P Natl Acad Sci USA 98:1095–1100

Takken W, Lindsay S (2019) Increased Threat of Urban Malaria from Anopheles stephensi Mosquitoes, Africa. Emerg Infect Dis 25:1431–1433

Taylor LR (1974) Insect Migration, Flight Periodicity and Boundary-Layer. J Anim Ecol 43:225-&

Tsoar A, Shohami D, Nathan R (2011) A Movement Ecology Approach to Study Seed Dispersal and Plant Invasion: An Overview and Application of Seed Dispersal by Fruit Bats. Fifty Years of Invasion Ecology: The Legacy of Charles Elton:103–119

Turner RM, Brockerhoff EG, Bertelsmeier C, et al. (2021) Worldwide border interceptions provide a window into human-mediated global insect movement. Ecol Appl 31

USDA (2023) 2023 USDA Plant Hardiness Zone Map. In: United States Department of Agriculture. https://planthardiness.ars.usda.gov/ Accessed: March 21, 2024 2024

WHO (2023) Invasive Vector Species: Anopheles stephensi. In: World Health Organization, Geneva, Switzerland (Web Page). https://apps.who.int/malaria/maps/threats/?theme=invasive&mapType=invasive%3A0&bounds=%5B%5B7.9910366929748875%2C-9.814831606483992%5D%2C%5B81.09046829208904%2C33.173143750325394%5D%5D&insecticideClass=PYRETHROIDS&insecticideTypes=&assayTypes=&synergistTypes=&species=&vectorSpecies=&surveyTypes=&deletionType=HRP2_PROPORTION_DELETION&plasmodiumSpecies=P._FALCIPARUM&drug=DRUG_AL&mmType=1&excludeLowerPatients=false&excludeLowerSamples=false&endemicity=false&countryMode=false&storyMode=false&storyModeStep=0&filterOpen=true&filtersMode=filters&years=1985%2C2023 Accessed: March 3, 2023 2023

Wu QL, Hu G, Tuan HA, et al. (2019) Migration patterns and winter population dynamics of rice planthoppers in Indochina: New perspectives from field surveys and atmospheric trajectories. Agr Forest Meteorol 265:99–109

Wu QL, Shen XJ, He LM, et al. (2021) Windborne migration routes of newly-emerged fall armyworm from Qinling Mountains-Huaihe River region, China. J Integr Agr 20:694–706

Yaro AS, Linton YM, Dao A, et al. (2022) Diversity, composition, altitude, and seasonality of high-altitude windborne migrating mosquitoes in the Sahel: Implications for disease transmission. Frontiers in Epidemiology 13 October 2022

Zahniser JN, Nartey R (2024) Curtara Insularis (Hemipera: Cicadellidae): New synonymy and correction of the type locality. Proceedings of the Entomological Society of Washington In Press

Zhou LY, Hoy CW, Miller SA, et al. (2003) Marking methods and field experiments to estimate aster leathopper dispersal rates. Environ Entomol 32:1177–1186

